# Evaluation of biodiversity in estuaries using environmental DNA metabarcoding

**DOI:** 10.1101/2020.03.22.997809

**Authors:** Hyojin Ahn, Manabu Kume, Yuki Terashima, Feng Ye, Satoshi Kameyama, Masaki Miya, Yoh Yamashita, Akihide Kasai

## Abstract

Biodiversity is an important parameter for the evaluation of the extant environmental conditions. Here, we used environmental DNA (eDNA) metabarcoding to investigate fish biodiversity in five different estuaries in Japan. Water samples for eDNA were collected from river mouths and adjacent coastal areas of two estuaries with high degrees of development (the Tama and Miya Rivers) and three estuaries with relatively low degrees of development (the Aka, Takatsu, and Sendai Rivers). A total of 182 fish species across 67 families were detected. Among them, 11 species occurred in all the rivers studied. Rare fishes including endangered species were successfully detected in rich natural rivers. Biodiversity was the highest in the Sendai River and lowest in the Tama River, reflecting the degree of human development along each river. Even though nutrient concentration was low in both the Aka and Sendai Rivers, the latter exhibited greater diversity, including many tropical or subtropical species, owing to its more southern location. Species composition detected by eDNA varied among rivers, reflecting the distribution and migration of fishes. Our results are in accordance with the ecology of each fish species and environmental conditions of each river, suggesting the potential of eDNA for non-invasive assessment of aquatic biodiversity.

## Introduction

As fisheries share common ecosystems and natural resources, concern has mounted over the impact of fishing on aquatic ecosystems [1]. To ensure sustainable fishery production, it is essential that organisms are reared in a balanced and healthy environment. At the same time, protecting rare and charismatic species has also gained importance [2]. One of the evaluation criteria for a balanced and healthy ecosystem is biodiversity.

Threats to biodiversity in aquatic ecosystems have been an issue for decades because of loss of productive habitats [3, 4]. Such environmental perturbations are caused mainly by human influences, through both direct damage to aquatic ecosystems and indirect pollution with sediments, excessive nutrients, and other chemicals. Terrestrial pollutants from agriculture, deforestation, and construction flow into coastal areas through the hydrologic system, mainly through rivers [5–7]. Therefore, humans affect first the estuaries and coastal areas, whose environmental conservation is indicated by the extent of biodiversity. Consequently, comprehensive monitoring of biodiversity is essential for conservation of ecosystems and sustainable fisheries production.

Although a number of studies on biodiversity have been reported [8, 9], most of them have focused on local areas of ecologic or economic importance to aquaculture [10], unique ecosystems (e.g., coral reefs, mangroves, tropical islands) [4, 6], and other services [11]. In contrast, biodiversity evaluations that include various regions at the same time have not been carried out, because traditional monitoring methods (observations and/or capture) require considerable financial and labor resources to cover a wide range of habitats [12, 13]. Also, particularly for rare and endangered species, monitoring using traditional methods can negatively affect the organisms and their habitat during the survey.

Here, we tested environmental DNA (eDNA) metabarcoding as a non-invasive and cost-effective method for monitoring the biodiversity of fishes [14] in multiple estuaries at a nation-wide scale. Environmental DNA, defined as genetic material released from organisms into the environment, has become a convenient tool for molecular biology and ecology over the past decade [15, 16]. By sampling soil, sediment, water, and ice, species can be detected even when they cannot be observed visually. This technique was first reported with regard to amphibians [17], followed by fish [18, 19], crustaceans [20], mammals [21], and plants [22]. In addition, combined with next-generation sequencing technology, eDNA enables the processing of massive DNA sequencing data for the identification of various taxa in multiple samples simultaneously, which is termed eDNA metabarcoding [23]. This method is not only practical for assessment of biodiversity, but is also useful to for detection of non-invasive alien, rare, and endangered species while performing a diversity survey [16, 24, 25]. We used specially designed universal primers covering 880 fish species belonging to 51 orders, 242 families, and 623 genera (MiFish-U), and 160 elasmobranch species belonging to 12 orders, 39 families, and 77 genera (MiFish-E) for the metabarcoding process [26].

Five rivers, indicative of different geographical features and human impact on biodiversity, were selected for this study. As Japan stretches extensively from north to south, the latitude of the target rivers varied from 31.85°N to 38.85°N (Fig 1a). The catchment area of the rivers showed considerable variation from natural forest to a megacity. We hypothesized that fish diversity detected from the eDNA survey would reflect those environmental characteristics.

**Fig 1.**
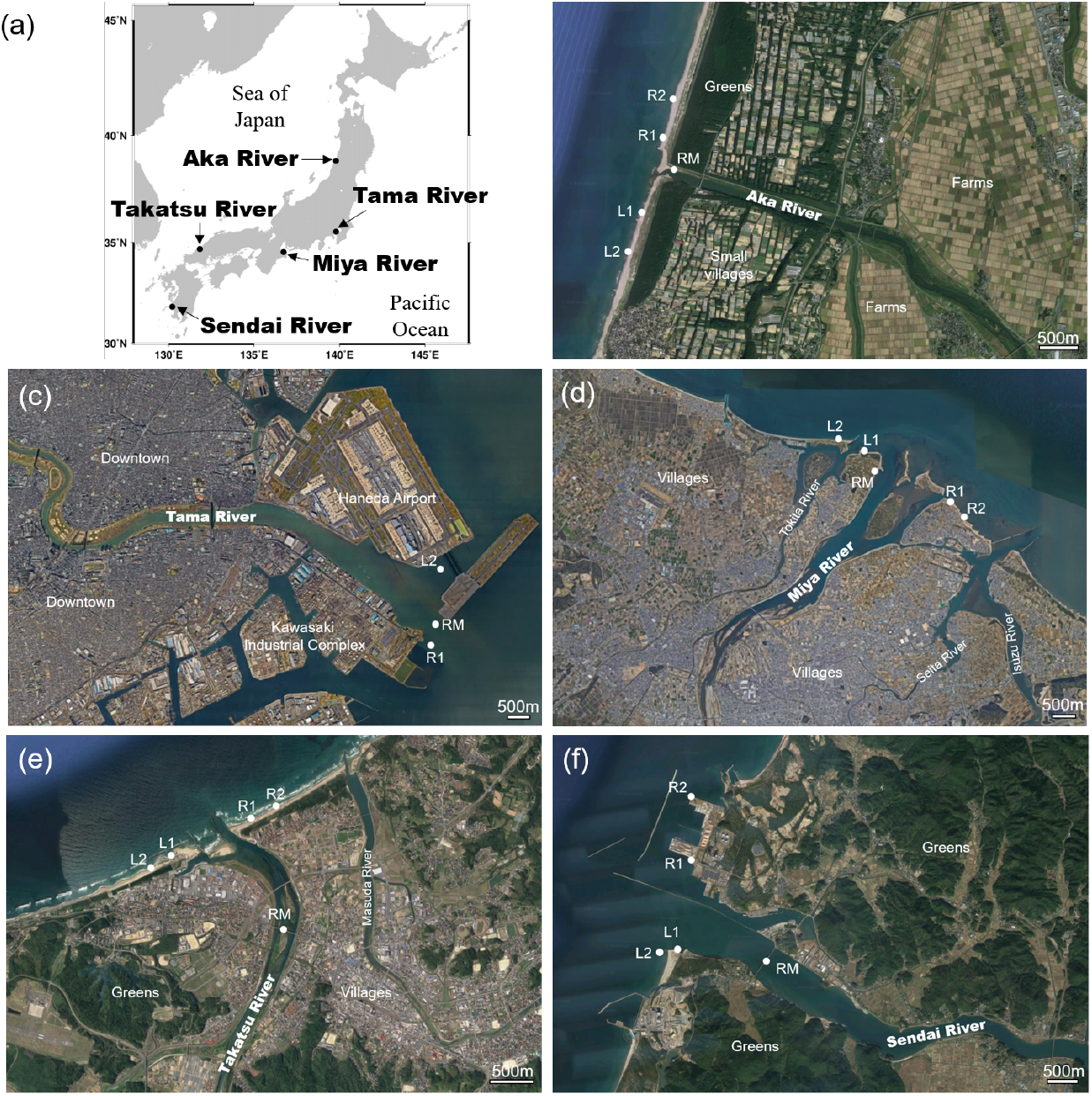
Sampling stations. Location of (a) the five rivers surveyed in this study. Maps showing the location of sampling stations RM (river mouth), L1 (left 500 m), L2 (left 1 km), R1 (right 500 m), and R2 (right 1 km) of (b) Aka River, (c) Tama River, (d) Miya River, (e) Takatsu River, and (f) Sendai River. The satellite photos from (b) to (f) were provided by Google Maps (2019 Google, TerraMetrics, Data SIO, NOAA, U.S. Navy, NGA, GEBCO). Scale bar = 500 m.

## Materials and methods

### Water sampling

Five rivers (Aka, Tama, Miya, Takatsu, and Sendai) with different geographical features and degrees of urbanization were selected. The water at five stations (at the river mouth, and approximately 500 m and 1 km along the coast on both the left and right sides of the river mouth) was sampled (Fig 1) in summer (June or July) 2018. At the river mouth, the water was sampled twice (at high and low tides), and therefore, there was a total of six samples collected from each estuary. For the Tama River, water samples were taken from a boat because the estuary is located between Haneda Airport and the Kawasaki industrial complex, and we could not reach the area from the shore. Moreover, because the airport restricts access to any type of boat near the runway, we could only collect samples from one station (at about 1 km from the river mouth) on each side of the Tama River estuary and collected four samples instead of six samples (Table 1).

**Table 1.** Environmental conditions of sampling stations. HT: river mouth at high tide; LT: river mouth at low tide; L1: left 500 m; L2: left 1 km; R1: right 500 m; R2: right 1 km.

| Aka River (17th, July) | HT | LT | L1 | L2 | R1 | R2 |
| --- | --- | --- | --- | --- | --- | --- |
| Water temp (°C) | 24.1 | 26 | 26 | 26 | 25.4 | 25.5 |
| Salinity | 6.4 | 7.7 | 18.8 | 29 | 20.1 | 29.7 |
| Filtered water (mL) | 200 | 200 | 600 | 600 | 600 | 600 |
| Tama River (29th, June) | HT | LT | L1 | L2 | R1 | R2 |
| Water temp (°C) | 22.3 | 26.6 | - | 24.7 | 23.5 | - |
| Salinity | 29.1 | 22.2 | - | 24.6 | 27.9 | - |
| Filtered water (mL) | 400 | 400 | - | 400 | 400 | - |
| Miya River (7th, June) | HT | LT | L1 | L2 | R1 | R2 |
| Water temp (°C) | 23 | 22.5 | 22.2 | 22.6 | 23.1 | 24.7 |
| Salinity | 7.45 | 10.4 | 25.53 | 23.39 | 20.64 | 21.92 |
| Filtered water (mL) | 700 | 600 | 200 | 500 | 600 | 600 |
| Takatsu River (16th, July) | HT | LT | L1 | L2 | R1 | R2 |
| Water temp (°C) | 26.7 | 24.1 | 29 | 29 | 27 | 27 |
| Salinity | 0.1 | 0.1 | 22.4 | 23.9 | 15.1 | 15.5 |
| Filtered water (mL) | 500 | 600 | 500 | 500 | 500 | 600 |
| Sendai River (27th, July) | HT | LT | L1 | L2 | R1 | R2 |
| Water temp (°C) | 28.7 | 30 | 29.7 | 29.5 | 29.1 | 29.3 |
| Salinity | 22.2 | 6.8 | 14.2 | 14.6 | 28.3 | 29.6 |
| Filtered water (mL) | 500 | 500 | 500 | 600 | 1000 | 1000 |

All sampling and filtering equipment was cleaned with 10% commercial bleach solution. The surface water at each station was sampled by a bucket and immediately filtered using a 0.45-μm polyethersulfone membrane Sterivex filter unit (Merck Millipore, Billerica, MA, USA) and immersed in 1.6 mL RNAlater Stabilization Solution (Thermo Fisher Scientific, Waltham, MA, USA). Water temperature and salinity were measured during sampling. The volume of water samples varied from 200 to 1000 mL depending on turbidity (Table 1). We assumed that variation in sample volume did not affect diversity as we confirmed no correlation between the volume and eDNA concentration (r^2^ = 0.045). As a negative control, 500 mL of pure water was filtered at each river. Filter units were frozen at −30°C until DNA extraction.

### eDNA extraction

Total DNA was extracted from the Sterivex filter units using a DNeasy Blood and Tissue Kit (Qiagen, Hilden, Germany), following the procedure described by Miya et al. [27] and the manufacturer’s protocol with minor modifications. After removing RNAlater by centrifugation (4,000 × *g* for 2 min), the filter unit was rinsed with sterilized distilled water. For the lysis of eDNA attached to the membrane, proteinase K (20 μL) and lysis buffer AL (200 μL) were applied to the filter unit and incubated inside a 56°C preheated oven for about 20 min. The roller was turned on to enable even collection of DNA from the membrane. After the incubation, the spin column was centrifuged at 4,000 × *g* for 2 min to collect DNA, to which 200 μL of absolute ethanol was then added and mixed well. The resulting solution was transferred to a spin column, centrifuged (6,000 × *g* for 1 min), and then purified twice using wash buffer (AW1 and AW2). After the purification steps, DNA was eluted with the elution buffer (110 μL) provided in the kit. Extracted DNA was stored in a LoBind tube at −30°C.

### Library preparation and sequencing

Samples were sent to the Kazusa DNA Research Institute (Chiba, Japan) for paired-end library preparation and next-generation sequencing (MiSeq) as detailed by Miya et al. [26].

A two-step PCR for paired-end library preparation was employed in the MiSeq platform (Illumina, San Diego, CA, USA). For the first-round PCR (1st PCR), a mixture of the following four primers was used: MiFish-U-forward (5’–ACA CTC TTT CCC TAC ACG CTC TTC CGA TCT NNN GTC GGT AAA ACT CGT GCC AGC–3’), MiFish-U-reverse (5’–GTG ACT GGA GTT CAG ACG TGT GCT CTT CCG ATC TNN NNN NCA TAG TGG GGT ATC TAA TCC CAG TTT G–3’), MiFish-E-forward-v2 (5’–ACA CTC TTT CCC TAC ACG CTC TTC CGA TCT NNN RGT TGG TAA ATC TCG TGC CAG C–3’), and MiFish-E-reverse-v2 (5’–GTG ACT GGA GTT CAG ACG TGT GCT CTT CCG ATC TNN NNN NGC ATA GTG GGG TAT CTA ATC CTA GTT TG–3’). These primer pairs amplified a hypervariable region of the mitochondrial 12S rRNA gene (*ca*. 172 bp; hereafter called “MiFish sequence”) and appended primer-binding sites (5’ ends of the sequences before six Ns) for sequencing at both ends of the amplicon. The six random bases (Ns) were used in the middle of these primers to enhance cluster separation in the flow cells during initial base call calibrations of the MiSeq platform.

The 1st PCR was carried out with 35 cycles of a 12-μL reaction volume containing 6.0 μL 2 × KAPA HiFi HotStart ReadyMix (KAPA Biosystems, Wilmington, MA, USA), 2.8◻μL of a mixture of the four MiFish primers in equal volumes (U/E forward and reverse primers; 5 μM), 1.2 μL sterile distilled water, and 2.0 μL eDNA template (a mixture of the duplicated eDNA extracts in equal volumes). To minimize PCR dropouts during the 1st PCR, eight replications were performed for the same eDNA template using a strip of eight tubes (0.2 μL). After an initial 3 min denaturation at 95°C, the thermal cycle profile (35 cycles) was as follows: denaturation at 98°C for 20 s, annealing at 65°C for 15 s, and extension at 72°C for 15 s. There was a final extension at 72°C for 5 min. The 1st PCR blanks were prepared during this process in addition to negative controls for each river.

After completion of the 1st PCR, equal volumes of the PCR products from the eight replications were pooled in a single 1.5-mL tube and purified using a GeneRead Size Selection kit (Qiagen) following the manufacturer’s protocol for the GeneRead DNA Library Prep I Kit. Accordingly, column purification was performed twice to completely remove adapter dimers and monomers. Subsequently, the purified target products (*ca*. 300 bp) were quantified using TapeStation D1000 (Agilent Technologies, Tokyo, Japan), after diluting them to 0.1 ng μL^−1^ with Milli Q water. The diluted products were employed as templates for the second-round PCR (2nd PCR).

For the 2nd PCR, the following two primers were used to append dual-index sequences (eight nucleotides indicated by Xs) and flow cell-binding sites for the MiSeq platform (5’ ends of the sequences before eight Xs): 2nd-PCR-forward (5’–AAT GAT ACG GCG ACC ACC GAG ATC TAC ACX XXX XXX XAC ACT CTT TCC CTA CAC GAC GCT CTT CCG ATC T–3’) and 2nd-PCR-reverse (5’–CAA GCA GAA GAC GGC ATA CGA GAT XXX XXX XXG TGA CTG GAG TTC AGA CGT GTG CTC TTC CGA TCT–3’).

The 2nd PCR was carried out with 10 cycles in a 15-μL reaction volume containing 7.5 μL 2 × KAPA HiFi HotStart ReadyMix, 0.9 μL of each primer (5 μM), 3.9 μL sterile distilled water, and 1.9 μL template (0.1 ng μL^−1^ except for the three blanks). After an initial 3 min denaturation at 95°C, the thermal cycle profile (10 cycles) was as follows: denaturation at 98°C for 20 s, combined annealing and extension at 72°C for 15 s. There was a final extension at 72°C for 5 min. The blank for the 2nd PCR was prepared during this process as well as to monitor any contamination.

All dual-indexed libraries were pooled in equal volumes into a 1.5-mL tube. Then, the pooled dual-indexed library was separated on a 2% E-Gel Size Select agarose gel (Life Technologies, Carlsbad, CA, USA) and the target amplicons (*ca*. 370 bp) were retrieved from the recovery wells using a micropipette. The concentration of the size-selected libraries was measured using a Qubit dsDNA HS assay kit and a Qubit fluorometer (Life Technologies). The libraries were diluted to 12.0 pM with HT1 buffer (Illumina) and sequenced on the MiSeq platform using a MiSeq v2 Reagent Kit for 2 × 150 bp PE (Illumina) following the manufacturer’s protocol.

### Data preprocessing and taxonomic assignment

Data preprocessing and analysis of MiSeq raw reads were performed with a specially developed pipeline (MiFish ver. 2.3) from four runs using USEARCH v10.0.240 [28]. The following steps (summarized in S1 Table) were applied: (1) Forward (R1) and reverse (R2) reads were merged by aligning them with the *fastq_mergepairs* command. During this process, the following reads were discarded: low-quality tail reads with a cut-off threshold set at a quality (Phred) score of 2, reads that were too short (<100 bp) after tail trimming, and paired reads with multiple differences (>5 positions) in the aligned region (*ca*. 65 bp). (2) Primer sequences were removed from merged reads using the *fastx_truncate* command. (3) Reads without primer sequences underwent quality filtering using the *fastq_filter* command to remove low-quality reads with an expected error rate >1% and reads that were too short (<120 bp). (4) Preprocessed reads were dereplicated using the *fastx_uniques* command and all singletons, doubletons, and tripletons were removed from subsequent analysis as recommended [28]. (5) Dereplicated reads were denoised using the *unoise3* command to generate amplicon sequence variants (ASVs) without any putatively chimeric and erroneous sequences [29]. (6) Finally, ASVs were subjected to taxonomic assignments of species names (metabarcoding operational taxonomic units; MOTUs) using the *usearch_global* command with sequence identity >98.5% to the reference sequences and a query coverage ≥90% (two nucleotide differences allowed). ASVs with sequence identities of 80–98.5% were tentatively assigned “U98.5” labels before the corresponding species name with the highest identity (*e.g.*, U98.5_*Pagrus*_*major*) and they were subjected to clustering at the 0.985 level using the *cluster_smallmem* command. In an incomplete reference database, this clustering step enables the detection of multiple MOTUs under an identical species name. We annotated such multiple MOTUs with “gotu1, 2, 3…” and tabulated all of these outputs (MOTUs plus U98.5_MOTUs) with read abundances. We excluded ASVs with sequence identities <80% (saved as “no_hit”) from the above taxonomic assignments and downstream analyses because all of them were found to be non-fish organisms.

As a reference database, we assembled MiFish sequences from 5,691 fish species in Masaki Miya’s laboratory. In addition, we downloaded all fish whole mitochondrial genome and 12S rRNA gene sequences from NCBI as of 26 June 2017 and extracted MiFish sequences using a custom Perl script [26]. We combined the MiFish sequences from the two sources in a FASTA format and used the combined sequences as the custom reference database for taxonomic assignments. The final reference database consisted of 27,871 sequences from 7,555 species belonging to 2,612 genera and 464 families.

We refined the above automatic taxonomic assignments with reference to family-level phylogenies based on MiFish sequences from MOTUs, U98.5_MOTUs, and the reference sequences from those families. For each family, we assembled representative sequences (most abundant reads) from MOTUs and U98.5_MOTUs, and added all reference sequences from that family and an outgroup (a single sequence from a closely-related family) in a FASTA format. We subjected the FASTA file to multiple alignment using MAFFT [30] with a default set of parameters. We constructed a neighbor-joining tree with the aligned sequences in MEGA7 [31] using Kimura two-parameter distances. The distances were calculated using pairwise deletion of gaps and among-site rate variations modeled with gamma distributions (shape parameter = 1). We performed bootstrap resamplings (*n* = 100) to estimate statistical support for internal branches of the neighbor-joining tree and to root the tree with the outgroup.

We inspected a total of 82 family-level trees and revised the taxonomic assignments. For U98.5_MOTUs placed within a monophyletic group consisting of a single genus, we assigned that genus to unidentified MOTUs with “sp” plus sequential numbers (e.g., *Pagrus* sp1, sp2, sp3, …). For the remaining MOTUs ambiguously placed in the family-level tree, we assigned the family name with “sp” plus sequential numbers (e.g., Sparidae sp1, sp2, sp3, …).

All negative controls in sampling stations and PCR blanks were also analyzed using this pipeline. The reads corresponding to every fish detected in the negative control were deleted (S1 Table).

### Species verification

The species obtained by pipeline still needed to be verified because sequencing results comprised only a short region (170 bp) of 12S rRNA (Miya et al., 2015), and similar sequences might correspond to different species. Also, multiple species could be incorporated into a single species, and *vice versa*. We checked all species on the list with the original aligned sequences using the NCBI Basic Local Alignment Search Tool (http://blast.ncbi.nlm.nih.gov/Blast.cgi), and applied MEGA7 [31] to construct a phylogenetic tree for all stations characterized by occurrence of the same species. When several species shared the same or similar (>99%) aligned sequence, we confirmed the species identity by referring to species distribution reported by the IUCN (https://www.iucnredlist.org), FishBase (http://www.fishbase.de), illustrated books of Japanese fishes [32–34], and personal communications. For example, the Japanese black porgy (*Acanthopagrus schlegelii*) and the Okinawa seabream (*Acanthopagrus sivicolus*) have the same aligned sequence, but the Okinawa seabream cannot exist in the waters of any station from the present study. On the contrary, we combined two or more species that were considered to be local variations, even if their sequences differed substantially.

Species whose reads number amounted to <0.05% of total reads were deleted because they were potentially caused by contamination, as indicated by Andruszkiewicz et al. [35] with some modifications. If species that were obviously not expected in this area were detected, but represented commonly consumed food items, they were regarded as contamination and removed as well.

### Estimates of biodiversity

Even if fish biomass could be reportedly determined by eDNA [36], eDNA has been limited to certain species. Moreover, it has not been applied to metabarcoding because of species-specific amplification rates [37], environment-dependent degradation rates [18, 38], and PCR inhibition by environmental factors [12, 14]. Therefore, the estimate of biomass requires a complex model and the possible use of eDNA for this purpose needs to be verified. Biodiversity is sometimes calculated by functions such as ‘number of species’ and ‘biomass;’ however, as biomass information was not available in the present study, we considered ‘species richness’ as a proxy for ‘biodiversity.’

### Environmental data set

Data regarding nutrients were obtained from the Ministry of the Environment of Japan (http://water-repo.env.go.jp/water-repo/). We used the annual mean value of nutrient concentration combining total nitrogen (TN) and total phosphorus (TP) published in the Measurement Results of Water Quality in Public Waters in FY 2016 (Ministry of the Environment) as a water quality index of the river. The annual mean value is based on 6–12 measurements a year at each monitoring point. The monitoring points corresponding to the target watersheds (points using the TN and TP values) were the most downward points of each river.

The revetment rate was calculated by measuring the distance of artificially protected areas, such as concrete-sealed piers or concrete tetrapods, within a distance of 3 km on both sides of the river and shore from the river mouth, using Google Earth Pro (http://support.google.com/earth/answer/21995?hl=ja).

### Statistical analysis

To examine the effect of salinity or water temperature on the ratio of freshwater, brackish, or seawater species, we used general linear models (GLMs) with a negative binominal distribution and a log link function. To this end, we applied the *glm.nb* function in the *MASS* package. The number of freshwater, brackish, or seawater species in each sample was used as a response variable; salinity or water temperature were explanatory variables; and the total number of fish species represented an offset term. To verify the accuracy of the six models, the areas under the Receiver Operating Characteristic curves (AUCs) were calculated, using the *roc* function in the *pROC* package [39]. Accuracy was defined as low (AUC < 0.7), moderate (0.7 ≤ AUC < 0.9), and high (AUC ≥ 0.9) (Table 2).

**Table 2.**
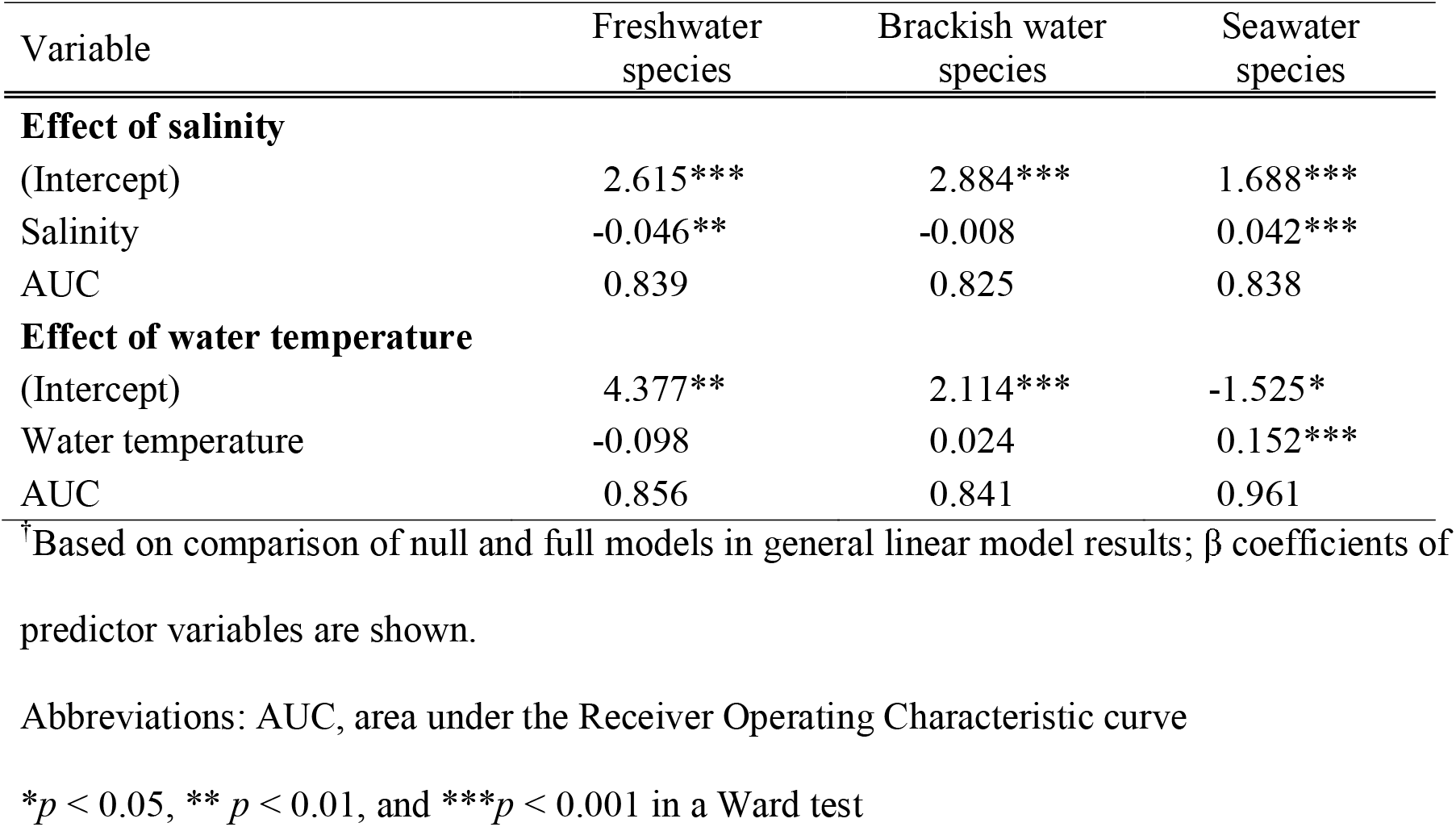
Summary of models^†^ used to assess the effect of each environmental factor on the rate of freshwater, brackish, or marine fish.

| Variable | Freshwater species | Brackish water species | Seawater species |
| --- | --- | --- | --- |
| **Effect of salinity** |  |  |  |
| (Intercept) | 2.615*** | 2.884*** | 1.688*** |
| Salinity | -0.046** | -0.008 | 0.042*** |
| AUC | 0.839 | 0.825 | 0.838 |
| **Effect of water temperature** |  |  |  |
| (Intercept) | 4.377** | 2.114*** | -1.525* |
| Water temperature | -0.098 | 0.024 | 0.152*** |
| AUC | 0.856 | 0.841 | 0.961 |

<sup>†</sup>Based on comparison of null and full models in general linear model results; $\beta$ coefficients of predictor variables are shown.
Abbreviations: AUC, area under the Receiver Operating Characteristic curve
\* $p < 0.05$ , \*\* $p < 0.01$ , and \*\*\* $p < 0.001$ in a Ward test

To examine the human impact on the number of fish species, we again applied the above GLMs using the *glm.nb* function in the *MASS* package. The number of species in each river was used as a response variable. We used data about TN, TP, and revetment rates as indicators of human impact. However, both TN and TP had a high variance inflation factor (VIF), which indicated high multicollinearity among these variables (VIF = 26.1 and 15.6 for TN and TP, respectively, VIF = 7.3 for revetment rate). After removal of TP, there was no multicollinearity between TN and revetment rate (VIF = 7.0), so we used TN and revetment rates as explanatory variables for our GLM analyses. These VIF values were calculated using the *vif* function in the *car* package [40]. The number of samples was used as an offset variable. For model selection among GLMs, we used the *dredge* function in the *MuMIn* package [41]. The best model was selected using Akaike’s information criterion (AIC), which stipulates that the best model for any candidate set applied to a given data set is that with the lowest AIC value. Following Burnham and Anderson [42], models with ΔAIC < 2 were assumed to be reasonable alternatives to the best model and thus were retained (Table 3).

**Table 3.**
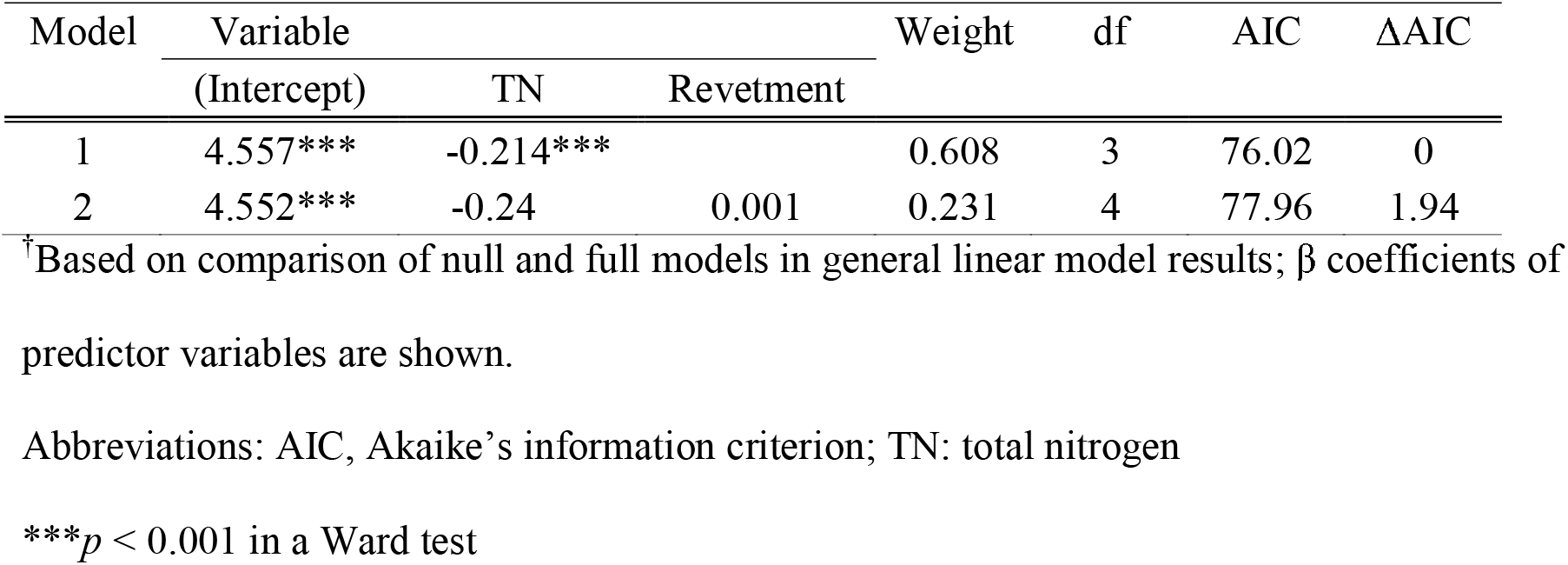
Summary of models with Δ AIC < 2^†^ used to assess the effect of human impact on the number of fish species.

| Model | Variable | | | Weight | df | AIC | $\Delta AIC$ |
| --- | --- | --- | --- | --- | --- | --- | --- |
|  | (Intercept) | TN | Revetment |  |  |  |  |
| 1 | 4.557*** | -0.214*** |  | 0.608 | 3 | 76.02 | 0 |
| 2 | 4.552*** | -0.24 | 0.001 | 0.231 | 4 | 77.96 | 1.94 |

<sup>†</sup>Based on comparison of null and full models in general linear model results; $\beta$ coefficients of predictor variables are shown.
Abbreviations: AIC, Akaike's information criterion; TN: total nitrogen
\*\*\* $p < 0.001$ in a Ward test

All statistical tests were carried out using R software ver. 3.5.2 [43].

## Results

### Species occurrence

A total of 182 species from 67 families were detected in the present eDNA survey (S2 Table). Most species (94) occurred in the Sendai River and fewest (25) in the Tama River; whereas the Aka, Miya, and Takatsu Rivers contributed with 64, 72, and 81 species, respectively (Fig 2a). Eleven species commonly observed in Japanese coastal areas (*Acanthogobius flavimanus*, *Acanthopagrus schlegelii*, *Cyprinus carpio*, *Engraulis japonicus*, *Girella punctata*, *Konosirus punctatus*, *Lateolabrax japonicus*, *Mugil cephalus*, *Parablennius yatabei*, *Platycephalus* sp. 2, and *Takifugu* spp.) were reported in all five rivers. Among them, the Japanese anchovy *E. japonicus* and dotted gizzard shad *K. punctatus* are commercially important; whereas the yellowfin goby *A. flavimanus*, blackhead seabream *A. schlegelii*, and *Platycephalus* sp. are popular for recreational fishing. Two salmonid species, *Oncorhynchus masou* and *Oncorhynchus mykiss*, known to inhabit colder and rural rivers were detected only in the Aka and Takatsu Rivers (S2 Table). Commercially and ecologically important fishes, such as the Japanese sardine *Sardinops melanostictus* and mackerel *Scomber* spp., were widely detected in all rivers except for the Sendai River.

**Fig 2.**
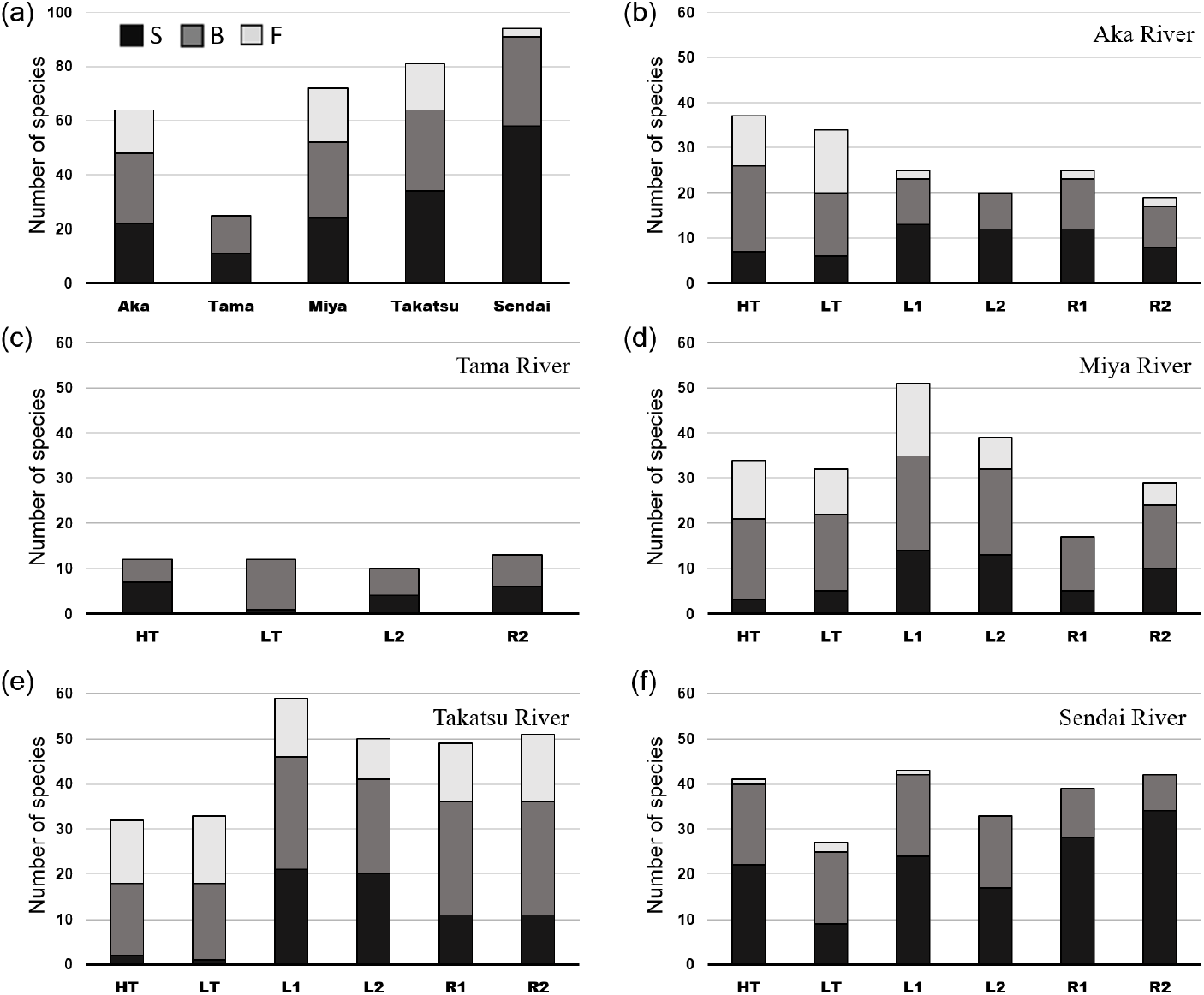
Species richness. Number of species present in (a) all five rivers and at each station, (b) Aka River, (c) Tama River, (d) Miya River, (e) Takatsu River, and (f) Sendai River. HT: river mouth at high tide; LT: river mouth at low tide; L1: left 500 m; L2: left 1 km; R1: right 500 m; R2: right 1 km; S: seawater species; B: brackish water species having a wide range of salinity tolerance including migrating fishes; F: freshwater species

*Cobitis takatsuensis*, *Hemitrygon akajei*, *Trachurus japonicus* (NT), *C. carpio*, *Hippocampus mohnikei* (VU), *Anguilla japonica*, and *Epinephelus akaara* (EN) are endangered according to the IUCN red list (https://www.iucnredlist.org). Moreover, *C. takatsuensis* and *A. japonica* are registered as endangered species at the EN level by the Ministry of the Environment of Japan (www. env.go.jp). An additional 11 species, detected by eDNA, including *Eutaeniichthys gilli*, *Gymnogobius castaneus*, *Misgurnus anguillicaudatus*, *Oncorhynchus masou*, *Sarcocheilichthys variegatus*, *Tanakia lanceolata* (NT), *Cottus kazika*, *Cottus reinii*, *Odontobutis hikimius* (VU), *Cottus pollux*, and *Gymnogobius scrobiculatus* (EN), are considered as endangered in Japan (http://ikilog.biodic.go.jp/Rdb/env).

### Habitat composition of each river

A detailed station-by-station analysis (Fig 2b–f) revealed that in the Tama River, freshwater species were not detected from all stations at the estuary (Fig 2c; S2 Table). Only a small proportion of freshwater species occurred at the river mouth and at the station 500 m left along the coast from the mouth of the Sendai River, while no freshwater species occurred at the other stations (Fig 2f). In the Aka River, freshwater species accounted for 30–40% of total species at the river mouth, but decreased quickly to fewer than 10% along both the left and right sides of the coast. In contrast, seawater species increased at stations in the coastal area (Fig 2b). Similar results were obtained for the Takatsu River, with the proportion of freshwater species decreasing and that of seawater species highly increasing in the coastal area (Fig 2e). A different result was observed regarding the number of species in the Aka and Takatsu Rivers (Fig 2b and 2e). More species were detected at the river mouth (37 species at high tide and 34 species at low tide) of the Aka River than in its surrounding coastal area (19– 25 species). In the Takatsu River, diversity was higher in the coastal area (49–59 species) than at the river mouth (32 at high tide and 33 species at low tide). In the Miya River, freshwater species decreased in the coastal area, except for the station at 500 m on the left side (Fig 2d). The number of species in the Sendai River decreased during low tide (27 species) compared to high tide (41 species) at the river mouth (Fig 2f). In the Tama River, species composition changed at the river mouth as the tide switched from high to low and seawater species decreased on the low tide, even though the total number of species (12 species) remained the same (Fig 2c). No distinguishable change was found between high and low tides at the river mouth of the other three rivers.

The best models examining the effect of salinity or water temperature on the ratio of freshwater, brackish, or seawater species could be obtained with relatively high accuracy (AUC = 0.825–0.961; Table 2). The proportion of freshwater species decreased as salinity increased (*p* < 0.01), whereas that of seawater species increased as salinity increased (*p* < 0.001) for all five rivers. In contrast, the proportion of brackish water fish was not affected by salinity. On the one hand, the proportion of seawater species increased at higher water temperatures (*p* < 0.001). On the other hand, water temperature had no significant effect on brackish and freshwater species (*p* > 0.05).

### Relationships between environmental factors and the number of species

Nutrient concentration (TN and TP) was highest in the Tama River, which flows through a mega city (Fig 1), and relatively low in the Aka and Takatsu Rivers, which flow through rural areas. A similar result was obtained regarding the revetment rate (Figs 1 and 3).

**Fig 3.**
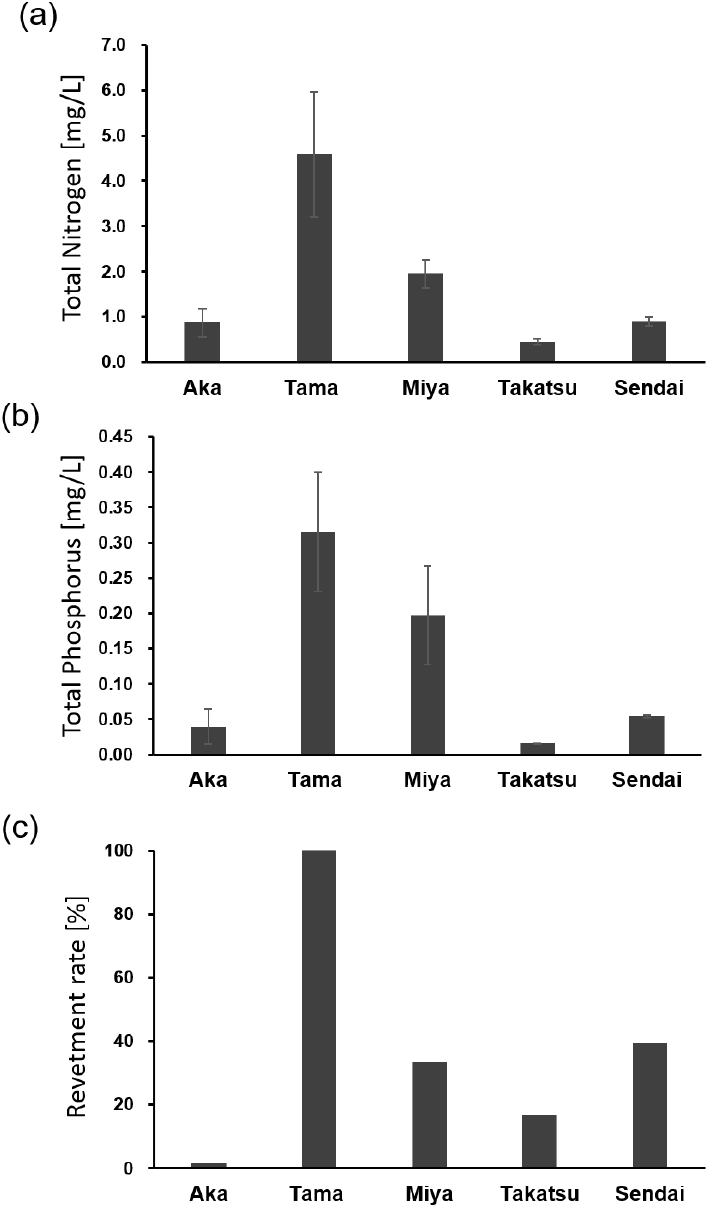
Human effects. Nutrients (mean ± SD) including (a) total nitrogen [mg/L] and (b) total phosphorus [mg/L] of the five rivers based on 2016 data obtained from the Ministry of the Environment, Japan (https://water-repo.env.go.jp/water-repo/). (c) Revetment rate [%] of the five rivers calculated using Google Earth Pro (2018 Google Image Landsat/Copernicus, US Dept of State Geographer Data SIO, NOAA, U. S. Navy, NGA, GEBCO). Bars show standard deviations.

Among the GLMs for evaluating the effect of human impact on the number of fish species, two models with ΔAIC < 2 were retained (Table 3). Both models included TN, whereby the number of species increased as TN decreased (*p* < 0.001). In the 2nd model, revetment was included but it had no significant effect (*p* > 0.05).

## Discussion

### Species composition

The 11 species detected in all five rivers are common in Japan, and some of them (e.g., *C. carpio* and *M. cephalus*) have a worldwide distribution [44, 45]. Some, such as *A. flavimanus* and *Takifugu* spp., can tolerate various environmental conditions [46, 47]. On the contrary, the endemic species *C. takatsuensis* was found only in a single habitat (i.e., the Takatsu River; S2 Table), confirming its known limited distribution [48]. This species is registered as an endangered species on the IUCN red list together with seven other species found in this study (https://www.iucnredlist.org). It is of particular importance that the endangered species were successfully detected by the eDNA survey as it is a non-intrusive method for both the environment and the subjects [14]. Therefore, eDNA could be applied not only for biodiversity research, but also to detect rare, endangered species [36]. Also, tropical to subtropical species (e.g., *Caranx ignobilis* [49]; *Spratelloides gracilis* [50]), only occurred in the Sendai River (S2 Table), which is located at the southernmost sampling station of the study. These results indicate that eDNA successfully reflects biological and geographical features.

Our eDNA analysis detected *K. punctatus* and *E. japonicus* in all estuaries (S2 Table). This result is consistent with the known distribution of these species; the former is distributed in estuaries from Tohoku southward and the latter in coastal areas across east Asia [33]. On the contrary, the herring *Clupea pallasii*, which is an important fisheries species in Japan, was not detected here as it is distributed in the north of Japan [33] and, hence, outside our study area. These results indicate that the eDNA survey adequately reflects coastal fish distribution. *S. melanostictus* was detected in all estuaries except the southernmost Sendai River estuary (S2 Table). This result is consistent with the ecology of *S. melanostictus*, which is known to migrate from south to north in summer. Our survey was conducted in June and July, and therefore it was not expected that *S. melanostictus* would be present in the Sendai River estuary, the southernmost observation point of this study. This finding indicates that a sequential eDNA survey can detect fish migration if a multipoint observation system is established, which would be useful especially for commercially important species.

### Environmental conditions and biodiversity

Biodiversity is closely related to the environmental conditions [10]. The results of GLMs showed that salinity affected the proportion of freshwater and seawater fishes, which varied among the five rivers. Specifically, no freshwater species eDNA samples were detected in the Tama River, which can be explained by the sampling stations being near the coast and salinity being over 20 (Table 1; Fig 1c). The proportion of seawater species accounted for more than 50% at high tide but decreased notably at low tide (Fig 2c). The Sendai River showed a very small proportion of freshwater species at the river mouth, which is relatively wide (>1 km), compared with the other four rivers (Figs 1f and 2f). It is believed that seawater easily enters into rivers with wide mouths, which causes freshwater from the river to disperse and dilute across the adjacent coastal areas. As a result, brackish and seawater species accounted for more than 90% of hits in this case.

Besides the width of rivers, tidal range is another factor with a strong influence on species composition. The tidal ranges are very small in the Sea of Japan [51], ranging from 6 cm for the Aka River to 55 cm for the Takatsu River, on the day of the sampling (www.jma.go.jp). In contrast, the tidal range of the Tama, Miya, and Sendai Rivers, which are located on the Pacific coast, was 167 cm, 67 cm, and 227 cm, respectively. Not surprisingly, salinity and number of species differed between high and low tides in the Sendai River (Table 1; Fig 2f). In the Tama River, the number of species did not differ between high and low tides; however, seawater species decreased at low tide (Fig 2c).

Species composition in the Aka River differed remarkably between the river mouth and coastal area; the proportion of freshwater species was about 30–40% at the river mouth but decreased to 8–10% in the coastal area, whereas seawater species increased from 18–19% at the river mouth to 42–60% in the coastal area. This pattern can also be explained by the width of the river mouth, which is very narrow (*ca*. 100 m) and thus affects species composition (Figs 1b and 2b). A similar trend was observed for the Takatsu River, which also has a narrow river mouth (<300 m); freshwater species decreased and seawater species increased in the coastal area. The proportion of seawater species was especially small at the river mouth of the Takatsu River, where water sampled from the bridge located about 1 km away from the river mouth had a salinity of 0.1 at both high and low tides (Table 1; Figs 1e and 2e). In fact, GLM analysis revealed that salinity had a significant effect on the proportion of freshwater and seawater species (Table 2).

Biodiversity was high at the river mouth of the Aka River, and in the coastal area of the Takatsu River (Fig 2b and e; S2 Table). As the number of species was almost identical at the river mouth of both rivers (34–37 species and 32–33 species, respectively), the observed change in biodiversity could be explained by two phenomena. First, as mentioned above, there are fewer freshwater species in the coastal area of the Aka River. Second, marine biodiversity is higher in the Takatsu River because it is located in the southern part of Japan and in general biodiversity increases toward lower latitudes [52]. GLM results supported the increase in number of seawater species when water temperature increased (Table 2).

Composition and number of species were less straightforward for the Miya River, reflecting its complex geography and environment (Fig 1d). For example, the number of species was highest at the station 500 m along the left of the river mouth (Fig 2d), which can be explained by the junction of two rivers, the Miya River and the Tokita River. However, the number of species was lowest at the station 500 m to the right of the river mouth, where no freshwater species were detected; the reason for this was not clear. The narrow river mouth beside the sampling station (R1) might prevent the flow of freshwater to the right side of the coast, but salinity was lower on the right side than on the left side, and some freshwater species were detected at the station 1 km to the right. One of the limitations and weaknesses of eDNA is the low amount of extracted DNA, which may not be enough for amplification and comprehensive species detection, as well as the presence of inhibitors such as humic acid, which might affect the results [14]. Therefore, although generally accurate, eDNA results might not always reflect all species present and other factors should be considered [53, 54].

Human activity exerts a large influence on the environment and biodiversity [7, 8]. Water quality is closely related to the biodiversity of aquatic animals [11]. Using nutrient concentrations (TN and TP) and revetment rate as indices of human activity and urbanization, we determined the impact of humans on biodiversity. GLM results indicated that TN significantly affected biodiversity, whereas the revetment rate had no effect (Table 3). The Tama River, which had the lowest biodiversity (Fig 2a), had the highest values for TN, TP, and revetment rates (Fig 3). The degree of urbanization of the Tama and Miya Rivers can be inferred not only from the concentration of nutrients and revetment rate but also from satellite images (Fig 1c and d). Even though the shoreline of the Sendai River has been extensively modified for flood control so that its revetment rate is now as high as for the Miya River, the surrounding area of the Sendai River has remained untouched and the nutrient concentration remains low (Figs 1 and 3). The Miya River showed relatively high biodiversity because of its location in the southern part of Japan along the Pacific coast, which is affected by the Kuroshio warm current. In comparison, even though it is located in the northern part of Japan, biodiversity was quite high in the Aka River (Fig 2a), which can be explained by the vastly pristine environment of the river (Fig 1b). This is an important result as it indicates that efforts to conserve the environment can also improve biodiversity. Both the Takatsu and Sendai Rivers showed high biodiversity with low human effect and geographical location (Figs 1, 2, and 3).

### Determination of sp. and spp

In cases whereby information was insufficient to determine the exact species, these were classified at the genus level (*Genus* sp.). If the species could not be confirmed either by sequencing or distribution, and more than two candidate species were possible, they were classified as *Genus* spp. Thus, *Carassius* spp. included the candidate species *C. auratus*, *C. cuvieri*, *C. gibelio*, and *C.* sp. CBM ZF 11717, all or only some of which could exist at the stations in the study. *Cypselurus* spp. might include *C. heterurus*, *C. hiraii*, and *C. opisthopus*. The genus *Cypselurus* was not supposed to be on the list as its habitat is far from the coast and *Cypselurus* species are found in several Japanese dishes. However, the sampling period covered the spawning season of this genus and this could influence the eDNA survey, so we decided to include it as a detected species. *Sebastes* spp. included *S. inermis*, *S. proriger*, *S. oblongus*, and *S. schlegelii*. *Hexagrammos* spp. included *H. agrammus*, *H. lagocephalus*, *H. otakii*, and *H. stelleri*. *Abudefduf* spp. included *A. sexfasciatus* and *A. vaigiensis*.

*Repomucenus* spp. included *R. beniteguri* and *R. omatipinnis*. *Acentrogobius* spp. included *A. pflaumii* and *A. virgatulus*. *Rhinogobius* spp. included *R. brunneus*, *R. flumineus*, *R. giurinus*, and *R.* sp. BF. *Tridentiger* spp. included *T. brevispinis* and *T. obscurus*. *Auxis* spp. included *A. rochei* and *A. thazard*. *Scomber* spp. included *S. australasicus* and *S. japonicus*. *Ostracion* spp. included *O. cubicus*, *O. immaculatus*, and *O. meleagris*. *Takifugu* spp. was probably *T. alboplumbeus* but could be also *T. flavipterus*, *T. pardalis*, *T. poecilonotus*, *T. porphyreus*, *T. rubripes*, *T. stictonotus*, and *T. xanthopterus* (S2 Table).

## Conclusion

The present study demonstrates that environmental DNA is a convenient tool for monitoring the distribution, migration, and diversity of fishes. By simply collecting 1 L of water, we successfully detected 182 species including commercially important species, covering a wide range of areas in a short period. The number and list of species detected by the eDNA method reflect the ecology of each fish and environmental conditions, such as eutrophication and temperature, in each river. We believe further development of the eDNA technique will offer an alternative method for accurate and non-invasive monitoring of aquatic life.

## Supporting information

Summary of data preprocessing steps and subsequent taxon assignment using pipeline analysis (MiFish ver. 2.3)

List of species detected at the sampling stations

## Acknowledgments

We are grateful to Dr. Komei Kadowaki and Mr. Shingo Takada for their help with fieldwork, Dr. Aya Yamazaki and Dr. Natsuko Kondo for guidance during molecular experiments and genetic analysis, Dr. Yoshiaki Kai and Dr. Yumi Henmi for confirming the species list, and Ms. Yuka Hayakawa for help with data analysis and arrangement.

## Supporting information

**S1 Table.** Summary of data preprocessing steps and subsequent taxon assignment using pipeline analysis (MiFish ver. 2.3)

**S2 Table.** List of species detected at the sampling stations. Plus (+) represents occurrence. HT: river mouth at high tide; LT: river mouth at low tide; L1: left 500 m; L2: left 1 km; R1: right 500 m; R2: right 1 km. ^†^: endangered species according to the IUCN (https://www.iucnredlist.org). ^‡^: endangered species according to the Ministry of the Environment of Japan (http://ikilog.biodic.go.jp/Rdb/env). ^†‡^: endangered species according to both classifications

