## Supplementary material for "Evaluation of biodiversity in estuaries using environmental DNA metabarcoding": Summary of data preprocessing steps and subsequent taxon assignment using pipeline analysis (MiFish ver. 2.3)

Table S1. Summary of data preprocessing steps and subsequent taxon assignments using pipeline analysis (MiFish ver. 2.3)

| Library | Raw read | Data preprocessing | | | Taxon assignment | |
|---|---|---|---|---|---|---|
| | | Merged | Quality filtered | Denoised | Fish | Non-fish |
| Aka (High tide) | 118,788 | 115,802 (97.5) | 114,784 (96.6) | 103,907 (87.5) | 103,835 (99.9) | 72 (0.1) |
| Aka (Low tide) | 103,740 | 101,039 (97.4) | 100,086 (96.5) | 90,209 (87.0) | 90,199 (100) | 10 (0.0) |
| Aka (Left 500 m) | 194,918 | 190,839 (97.9) | 189,593 (97.3) | 180,918 (92.8) | 180,914 (100) | 4 (0.0) |
| Aka (Left 1 km) | 173,568 | 166,071 (95.7) | 164,763 (94.9) | 153,667 (88.5) | 153,627 (100) | 40 (0.0) |
| Aka (Right 500 m) | 123,351 | 120,362 (97.6) | 119,344 (96.8) | 111,007 (90.0) | 110,988 (100) | 19 (0.0) |
| Aka (Right 1 km) | 128,843 | 125,802 (97.6) | 124,983 (97.0) | 118,721 (92.1) | 118,721 (100) | 0 (0.0) |
| Tama (High tide) | 79,490 | 71,151 (89.5) | 70,543 (88.7) | 65,652 (82.6) | 65,627 (100) | 25 (0.0) |
| Tama (Low tide) | 248,331 | 172,848 (69.6) | 170,955 (68.8) | 159,290 (64.1) | 159,234 (100) | 56 (0.0) |
| Tama (Left 1 km) | 284,947 | 164,667 (57.8) | 162,990 (57.2) | 151,642 (53.2) | 150,851 (99.5) | 791 (0.5) |
| Tama (Right 500 m) | 258,258 | 201,511 (78.0) | 199,299 (77.2) | 186,924 (72.4) | 186,742 (99.9) | 182 (0.1) |
| Miya (High tide) | 113,939 | 111,758 (98.1) | 110,988 (97.4) | 102,246 (89.7) | 102,242 (100) | 4 (0.0) |
| Miya (Low tide) | 95,202 | 93,526 (98.2) | 92,919 (97.6) | 86,238 (90.6) | 86,233 (100) | 5 (0.0) |
| Miya (Left 500 m) | 125,026 | 120,839 (96.7) | 119,879 (95.9) | 108,768 (87.0) | 107,356 (98.7) | 1,412 (1.3) |
| Miya (Left 1 km) | 83,342 | 81,639 (98.0) | 81,105 (97.3) | 74,467 (89.4) | 74,355 (99.8) | 112 (0.2) |
| Miya (Right 500 m) | 73,975 | 72,479 (98.0) | 71,979 (97.3) | 66,688 (90.2) | 66,688 (100) | 0 (0.0) |
| Miya (Right 1 km) | 98,725 | 96,859 (98.1) | 96,148 (97.4) | 89,019 (90.2) | 88,991 (100) | 28 (0.0) |
| Takatsu (High tide) | 168,953 | 164,080 (97.1) | 162,519 (96.2) | 149,184 (88.3) | 148,611 (99.6) | 573 (0.4) |
| Takatsu (Low tide) | 214,832 | 209,021 (97.3) | 206,950 (96.3) | 189,960 (88.4) | 188,096 (99.0) | 1,864 (1.0) |
| Takatsu (Left 500 m) | 194,703 | 186,526 (95.8) | 185,168 (95.1) | 169,508 (87.1) | 169,092 (99.8) | 416 (0.2) |
| Takatsu (Left 1 km) | 167,023 | 153,643 (92.0) | 152,448 (91.3) | 139,993 (83.8) | 139,633 (99.7) | 360 (0.3) |
| Takatsu (Right 500 m) | 202,839 | 197,007 (97.1) | 195,506 (96.4) | 179,825 (88.7) | 179,166 (99.6) | 659 (0.4) |
| Takatsu (Right 1 km) | 267,266 | 258,367 (96.7) | 256,276 (95.9) | 236,318 (88.4) | 234,913 (99.4) | 1,405 (0.6) |
| Sendai (High tide) | 83,224 | 81,469 (97.9) | 80,895 (97.2) | 73,927 (88.8) | 73,642 (99.6) | 285 (0.4) |
| Sendai (Low tide) | 170,223 | 166,437 (97.8) | 164,967 (96.9) | 155,201 (91.2) | 155,148 (100) | 53 (0.0) |
| Sendai (Left 500 m) | 147,759 | 144,440 (97.8) | 143,119 (96.9) | 132,680 (89.8) | 132,276 (99.7) | 404 (0.3) |
| Sendai (Left 1 km) | 177,691 | 171,791 (96.7) | 169,645 (95.5) | 157,985 (88.9) | 157,856 (99.9) | 129 (0.1) |
| Sendai (Right 500 m) | 151,445 | 148,697 (98.2) | 147,965 (97.7) | 140,021 (92.5) | 139,811 (99.9) | 210 (0.1) |
| Sendai (Right 1 km) | 192,176 | 186,251 (96.9) | 184,471 (96.0) | 171,052 (89.0) | 170,736 (99.8) | 316 (0.2) |
| Total | 4,442,577 | 4,074,921 (91.7) | 4,040,287 (90.9) | 3,745,017 (84.3) | 3,735,583 (99.7) | 9,434 (0.3) |
| Aka (Negative control) | 450 | 395 (87.8) | 387 (86.0) | 287 (63.8) | 280 (97.6) | 7 (2.4) |
| Tama (Negative control) | 14,815 | 14,338 (96.8) | 14,174 (95.7) | 12,811 (86.5) | 12,811 (100.0) | 0 (0.0) |
| Miya (Negative control) | 118 | 102 (86.4) | 98 (83.1) | 8 (6.8) | 8 (100.0) | 0 (0.0) |
| Takatsu (Negative control) | 870 | 804 (92.4) | 797 (91.6) | 594 (68.3) | 49 (8.2) | 545 (91.8) |
| Sendai (Negative control) | 1,020 | 915 (89.7) | 897 (87.9) | 690 (67.7) | 235 (34.1) | 455 (65.9) |
| 1st PCR blank-1 | 271 | 253 (93.4) | 248 (91.5) | 211 (77.9) | 0 0.0 | 211 (100) |
| 1st PCR blank-2 | 22 | 17 (77.3) | 16 (72.7) | 12 (54.6) | 0 0.0 | 12 (100) |
| 1st PCR blank-3 | 0 | 0 | 0 | 0 | 0 | 0 |
| 1st PCR blank-4 | 5 | 4 (80.0) | 4 (80.0) | 0 (0.0) | 0 (0.0) | 0 (0.0) |
| 1st PCR blank-5 | 1 | 1 (100) | 0 (0.0) | 0 (0.0) | 0 (0.0) | 0 (0.0) |
| 1st PCR blank-6 | 2 | 0 | 0 | 0 | 0 | 0 |
| 1st PCR blank-7 | 8 | 7 (87.5) | 7 (87.5) | 0 (0.0) | 0 (0.0) | 0 (0.0) |
| 1st PCR blank-8 | 18 | 17 (94.4) | 17 (94.4) | 6 (33.3) | 0 0.0 | 6 (100) |
| 1st PCR blank-1 | 0 | 0 | 0 | 0 | 0 | 0 |
| 1st PCR blank-2 | 0 | 0 | 0 | 0 | 0 | 0 |
| Total | 17,600 | 16,853 (95.8) | 16,645 (94.6) | 14,619 (83.1) | 13,383 (91.5) | 1,236 (8.5) |
