## Supplementary material for "Evaluation of biodiversity in estuaries using environmental DNA metabarcoding": List of species detected at the sampling stations

Table S2. List of species detected at the sampling stations. Plus (+) represents occurrence. HT: river mouth at high tide, LT: river mouth at low tide, L1: left 500 m, L2: left 1 km, R1: right 500 m, R2: right 1 km, †: endangered species according to the IUCN (https://www.iucnredlist.org), ‡: endangered species according to the Ministry of the Environment of Japan (http://ikilog.biodic.go.jp/Rdb/env), †‡: endangered species according to both classifications

| Family | Scientific Name | Aka | | | | | | Tama | | | | Miya | | | | | | Takatsu | | | | | | Sendai | | | | | |
|---|---|---|---|---|---|---|---|---|---|---|---|---|---|---|---|---|---|---|---|---|---|---|---|---|---|---|---|---|---|
| | | HT | LT | L1 | L2 | R1 | R2 | HT | LT | L2 | R1 | HT | LT | L1 | L2 | R1 | R2 | HT | LT | L1 | L2 | R1 | R2 | HT | LT | L1 | L2 | R1 | R2 |
| Dasyatidae | *Hemitrygon akajei* † | | | | | | | | | | | | + | + | | | | | | | | | | | | | | | |
| Anguillidae | *Anguilla japonica* †‡ | | | | | | | | | | | + | + | + | + | | | + | + | + | + | + | + | + | + | + | | | |
| Muraenidae | *Gymnothorax kidako* | | | | | | | | | | | | | | | | | | | | | | | | | | | + | |
| Engraulidae | *Engraulis japonicus* | | + | + | + | + | + | + | + | + | + | | + | | + | | | | | | | + | | + | + | + | + | + | + |
| Clupeidae | *Etrumeus micropus* | | | | | | | + | | | + | | | + | + | | | | | | | | | | | + | | | |
| | *Konosirus punctatus* | | + | + | + | + | + | + | + | + | + | + | + | + | + | | + | | | | | + | + | + | + | + | + | + | + |
| | *Sardinella zunasi* | | | | | | | | | | | | | + | | | | | | | | | | | | | | | |
| | *Sardinops melanostictus* | | | | | + | | + | | + | | | | | + | | + | | | | | | + | | | | | | |
| | *Spratelloides gracilis* | | | | | | | | | | | | | | | | | | | | | | | + | | + | + | + | + |
| Cyprinidae | *Acheilognathus rhombeus* | | | | | | | | | | | | | + | | | | | | | | | | | | | | | |
| | *Carassius* spp. | + | + | | | | | | | | | + | + | + | + | | + | + | + | + | + | + | + | | + | + | | | |
| | *Cyprinus carpio* † | + | + | | + | + | + | | | + | | + | + | + | + | | | + | + | + | + | + | + | + | + | + | | | |
| | *Gnathopogon elongatus elongatus* | + | + | | | | | | | | | + | + | + | + | | + | | | | | | | | | | | | |
| | *Hemibarbus labeo* | + | + | | | | | | | | | + | + | + | | | | | | | | | | | | | | | |
| | *Nipponocypris sieboldii* | | | | | | | | | | | | | + | | | | | | | | | | | | | | | |
| | *Nipponocypris temminckii* | | | | | | | | | | | + | | | | | | + | + | + | + | + | + | + | | | | | |
| | *Opsariichthys platypus* | + | + | | | | | | | | | + | + | + | + | | | + | + | + | + | + | + | | + | | | | |
| | *Phoxinus lagowskii steindachneri* | + | + | | | | | | | | | | | | | | | | | | | | | | | | | | |
| | *Phoxinus oxycephalus jouyi* | | + | | | | | | | | | | | | | | | + | + | | | + | + | | | | | | |
| | *Phoxinus* sp. | | | | + | + | | | | | | | | | | | | | | | | | | | | | | | |
| | *Pseudogobio esocinus esocinus* | + | + | | | | | | | | | + | + | + | + | | | + | + | + | + | + | + | | | | | | |
| | *Pseudorasbora parva* | + | | | | | | | | | | | | | | | | | | | | | | | | | | | |
| | *Pungtungia herzi* | | | | | | | | | | | | | | | | | + | + | + | + | + | + | | | | | | |
| | *Rhodeus ocellatus ocellatus* | | + | | | | | | | | | + | + | + | | | + | | | | | | | | | | | | |
| | *Sarcocheilichthys variegatus* ‡ | + | + | | | | | | | | | | | + | | | | | | | | | | | | | | | |
| | *Squalidus gracilis* | | | | | | | | | | | | | | | | | + | + | + | | + | + | | | | | | |
| | *Squalidus japonicus* | | | | | | | | | | | | | | + | | | | | | | | | | | | | | |
| | *Tanakia lanceolata* ‡ | | | | | | | | | | | + | + | + | + | | | | | | | | | | | | | | |
| | *Tanakia limbata* | | | | | | | | | | | + | + | + | | | + | + | + | + | | | + | | | | | | |
| | *Tribolodon brandtii maruta* | + | | | | | | + | + | | | | | | | | | | | | | | | | | | | | |
| | *Tribolodon hakonensis* | + | + | + | + | + | + | | + | | | + | + | + | + | | + | + | + | + | + | + | + | | | | | | |
| Cobitidae | *Cobitis minamorii tokaiensis* | | | | | | | | | | | + | | + | | | | | | | | | | | | | | | |
| | *Cobitis takatsuensis* †‡ | | | | | | | | | | | | | | | | | + | + | + | | + | + | | | | | | |
| | *Cobitis* sp. BIWAE type B | | | | | | | | | | | + | | | + | | | | | | | | | | | | | | |
| | *Misgurnus anguillicaudatus* ‡ | + | + | + | | + | + | | | | | | | + | | | | | + | + | | + | + | | | | | | |
| | *Misgurnus* sp. Clade A | | + | | | | | | | | | | | | | | | | | | | | | | | | | | |
| Balitoridae | *Barbatula* sp. | | | | | | | | | | | | | | | | + | | | | | | | | | | | | |
| | *Lefua echigonia* | | + | | | | | | | | | | | | | | | | | | | | | | | | | | |
| Siluridae | *Silurus asotus* | + | + | | | | | | | | | | | + | | | | + | + | + | + | + | + | | | | | | |
| Plotosidae | *Plotosus japonicus* | | | | | | | | | | | | | | | | | | | | + | | | | | | | | + |
| Bagridae | *Tachysurus nudiceps* | | | | | | | | | | | | | | | | | + | + | + | + | + | + | | | | | | |
| Osmeridae | *Plecoglossus altivelis altivelis* | | | | | | | | | | | + | + | + | | | | + | + | + | + | + | + | | | | | | |
| | *Salangichthys ishikawae* | | | | | | | | | | | | | + | | | | | | | | | | | | | | | |
| Salmonidae | *Oncorhynchus masou* ‡ | + | + | | | | | | | | | | | | | | | + | + | + | + | + | + | | | | | | |
| | *Oncorhynchus mykiss* | + | | | | | | | | | | | | | | | | | | | | | | | | | | | |
| Mugilidae | *Liza affinis* | | | | | | | | | | | | | | | | | | | | | | | + | + | + | | | |
| | *Liza haematocheila* | + | + | | | | | | | | | | | | | | | | + | | | | | | | | | | |
| | *Mugil cephalus* | + | + | | | + | + | + | + | + | + | + | + | + | + | + | + | + | + | + | + | + | + | + | + | + | + | + | + |
| Atherinidae | *Atherion elymus* | | | | | | | | | | | | | | | | | | | | | | | | | | | + | + |
| | *Hypoatherina tsurugae* | | | | | | | | | | | | | | | | | | | | | | | | | | | + | |
| | *Hypoatherina valenciennei* | | | | | | | | | | | | | + | + | | | | | | | | | | | | | | |
| Exocoetidae | *Cypselurus* spp. | | | + | + | + | + | | | | | | | | | | | | | + | + | | | | | | | | |
| Hemiramphidae | *Hyporhamphus sajori* | | | + | + | | | | | | | + | + | + | + | | + | | | + | + | + | + | | | | | | |
| Belonidae | *Ablennes hians* | + | + | | | | | | | | | | | | | | | | | + | + | | | | | | | + | |
| | *Strongylura anastomella* | + | + | + | + | + | + | | | | | | | | | + | + | | | + | + | + | + | + | + | + | | | |
| | *Tylosurus crocodilus* | | | | | | | | | | | | | | | | | | | | | | | + | + | + | + | | |
| Syngnathidae | *Hippocampus mohnikei* † | | | | | | | | | | | | | + | | | | | | | | | | | | | | | |
| Scorpaenidae | *Sebastes* spp. | | | + | | | | | | | | | + | | + | | | | | | | | | | | | | | |
| | *Sebastiscus marmoratus* | | | | | | | | | | | | | | + | | | | | + | | | | | | + | | + | + |
| Platycephalidae | *Platycephalus* sp. 2 | | | | | | + | | + | | | | | + | + | + | + | | | + | | + | + | | | + | | | |
| Hexagrammidae | *Hexagrammos* spp. | | | + | | + | | | | | | | | | | | | | | + | + | | | | | | | | |
| Cottidae | *Cottus kazika* ‡ | + | | | | | | | | | | + | | | | + | | + | + | + | + | + | + | | | | | | |
| | *Cottus pollux* ‡ | + | + | | | + | | | | | | | | | | | | + | + | + | + | + | + | | | | | | |
| | *Cottus reinii* ‡ | | | | | | | | | | | + | + | + | | | | | | | | | | | | | | | |
| Moronidae | *Lateolabrax japonicus* | + | + | + | + | + | + | + | + | + | + | + | + | + | + | + | + | + | + | + | + | + | + | + | + | + | + | + | |

Table S2. Continued.

| Family | Scientific Name | Aka | | | | | | Tama | | | | Miya | | | | | | Takatsu | | | | | | Sendai | | | | | |
|---|---|---|---|---|---|---|---|---|---|---|---|---|---|---|---|---|---|---|---|---|---|---|---|---|---|---|---|---|---|
| | | HT | LT | L1 | L2 | R1 | R2 | HT | LT | L2 | R1 | HT | LT | L1 | L2 | R1 | R2 | HT | LT | L1 | L2 | R1 | R2 | HT | LT | L1 | L2 | R1 | R2 |
| Moronidae | *Lateolabrax latus* | | | | | | | | | | | + | + | + | + | + | + | | | + | + | + | + | + | + | + | + | + | + |
| | *Lateolabrax maculatus* | | | | | | | | | | | | | | | | | | | | | | | | + | + | | | |
| Serranidae | *Diploprion bifasciatum* | | | | | | | | | | | | | | | | | | | + | | | | | | | | | |
| | *Epinephelus akaara* † | | | | | | | | | | | | | | | | | | | + | | | + | | | | | | |
| | *Epinephelus awoara* | | | | | | | | | | | | | | | | | | | | | | | + | | | | | |
| Centrarchidae | *Micropterus salmoides* | | | | | | | | | | | | | | | | | + | | | | | | | | | | | |
| Apogonidae | *Apogon coccineus* | | | | | | | | | | | | | | | | | | | | | | | | | | | + | |
| | *Apogon unicolor* | | | | | | | | | | | | | | | | | | | | | | | + | | | | | |
| | *Ostorhinchus doederleini* | | | | | | | | | | | | | | | | | | | | | | | | | | | + | + |
| | *Ostorhinchus notatus* | | | | | | | | | | | | | | | | | | | | | | | | | | | + | + |
| | *Ostorhinchus semilineatus* | | | | | | | | | | | | | | | | | | | | + | | | | | | | + | + |
| Sillaginidae | *Sillago japonica* | + | + | + | + | + | + | | | | | | | + | + | | + | + | | + | + | + | + | + | + | + | + | + | + |
| Carangidae | *Caranx ignobilis* | | | | | | | | | | | | | | | | | | | | | | | + | | | | | |
| | *Decapterus maruadsi* | | | | | | | | | | | | | | | | | | | + | | | | | | | | | |
| | *Kaiwarinus equula* | | | | | | | | | | | | | | | | | | | | + | | | | | | | | |
| | *Seriola dumerili* | | | | | | | | | | | | | | | | | | | | | | | + | | | | | |
| | *Seriola quinqueradiata* | + | | | | | | | | | | | | | | | | | | + | | | | | | | | | |
| | *Trachurus japonicus* † | + | | + | + | + | + | | | | | | | | | | | | | + | + | + | + | + | + | + | + | + | + |
| Leiognathidae | *Equulites rivulatus* | | | | | | | | | | | | | | | | | | | | | | | | | + | | + | |
| | *Nuchequula nuchalis* | + | + | + | | + | + | | | | | | | | + | | | | | | | | | | | | | | |
| Lutjanidae | *Lutjanus ophuysenii* | | | | | | | | | | | | | | | | | | | | + | | | | | | | | |
| Gerreidae | *Gerres equulus* | | | | | | | | | | | | | | | | | | | | | | | | | | | | + |
| Haemulidae | *Parapristipoma trilineatum* | | | | | | | | | | | | | + | | | | | | + | | + | | | | | + | | |
| | *Plectorhinchus cinctus* | | | | | | | | | | | | | | | | | | | | | | | + | + | + | + | | |
| Sparidae | *Acanthopagrus latus* | | | | | | | | | + | + | + | | | | | | | | | | | | + | + | + | + | | |
| | *Acanthopagrus schlegelii* | + | | + | + | + | + | + | + | + | + | + | + | + | + | + | + | + | + | + | + | + | + | + | + | + | + | + | + |
| | *Pagrus major* | | + | | + | | + | | | | | | | | | | | | | | | + | | + | | + | + | + | + |
| | *Rhabdosargus sarba* | | | | | | | | | | | | + | | + | | | | | | | | | | | | | + | |
| Polynemidae | *Polydactylus plebeius* | | | | | | | | | | | | | | | | | | | | | | | | | | + | | |
| Sciaenidae | *Nibea mitsukurii* | | | | | | | | | | | | | | | | | | | | | | | + | | + | | | |
| | *Pennahia argentata* | | | | + | + | | | | | + | | | | | | | | | | | | | | | | | | |
| Mullidae | *Parupeneus ciliatus* | | | | | | | | | | | | | | | | | | | | | | | | | | + | | |
| Pempheridae | *Pempheris* sp. | | | | | | | | | | | | | | | | | | | | | | | | | | | + | + |
| Kyphosidae | *Girella leonina* | | | | | | | | | | | | | | | | | | | | | | | | | + | + | + | + |
| | *Girella punctata* | + | | + | | + | | + | | | + | + | | | | | + | | | + | + | + | | + | | + | + | + | + |
| | *Kyphosus bigibbus* | | | | | | | | | | | | | | | | | | | | | | | | | | | | + |
| | *Microcanthus strigatus* | | | | | | | | | | | | | | | | | | | | | | | | | | + | | |
| Chaetodontidae | *Chaetodon auripes* | | | | | | | | | | | | | | | | | | | | | | | | | | | + | + |
| Teraponidae | *Rhynchopelates oxyrhynchus* | | | | | | | | | | | | | | | | | | | | + | + | + | + | + | + | + | | + |
| Oplegnathidae | *Oplegnathus fasciatus* | | | | | | | | | | | | | | | | | | | | + | | | | | | | | |
| Cheilodactylidae | *Goniistius* sp. | | | | | | | | | | | | | | | | | | | | + | | | | | | | | |
| Embiotocidae | *Ditrema temminckii* | | | | | | | | | | + | | | | | | | | | | | | + | | | | | | |
| Pomacentridae | *Abudefduf* spp. | | | | | | | | | | | | | | | | | | | | | | | | | | | + | + |
| | *Pomacentrus coelestis* | | | | | | | | | | | | | | | | | | | | | | | | | | | | + |
| | *Pomacentrus nagasakiensis* | | | | | | | | | | | | | | | | | | | | | | | | | | | | + |
| Labridae | *Choerodon azurio* | | | | | | | | | | | | | | | | | | | | + | | | | | | | | |
| | *Halichoeres poecilopterus* | | | | | | | | | | | | | | | | | | | + | + | | | | | | | | |
| | *Halichoeres tenuispinis* | | | | | | | | | | | | | | | | | | | | | | | | | | + | | + |
| | *Pseudolabrus eoethinus* | | | | | | | | | | | | | | | | | | | | | | | | | | | + | + |
| | *Pseudolabrus sieboldi* | | | | | | | | | | | | | | | | | | | | | | | | | | | | + |
| | *Stethojulis interrupta* | | | | | | | | | | | | | | | | | | | | | | | | | | | + | |
| | *Thalassoma lunare* | | | | | | | | | | | | | | | | | | | | | | | | | | | | + |
| | *Thalassoma cupido* | | | | | | | | | | | | | | | | | | | | | | | | | | | + | + |
| Stichaeidae | *Dictyosoma burgeri* form b | | | | | | | | | | | | | | | | | | | + | | + | | | | + | | | |
| Pholidae | *Pholis nebulosa* | | | | | | | | | | | | | | + | | | | | | | | | + | | | | | |
| Tripterygiidae | *Enneapterygius etheostomus* | | | + | | | | | | | | | | | | | | | | + | + | | | | | | | + | |
| | *Enneapterygius* sp. | | | | | | | | | | | | | | | | | | | | | | | | | | | | + |
| Blenniidae | *Entomacrodus stellifer* | | | | | | | | | | | | | | | | | | | + | + | | | + | | + | + | | + |
| | *Istiblennius enosimae* | | | | | | | | | | | | | | | | | | | | | | | | | | + | + | + |
| | *Omobranchus elegans* | | | + | | | | | | | | | | + | | + | | | | | + | | | | | | | | |
| | *Omobranchus fasciolatoceps* | | | | | | | | | | | | | | | | + | | | | | | | + | | + | | | |
| | *Omobranchus loxozonus* | | | | | | | | | | | | | | | | | | | | | | | | | | | | + |
| | *Omobranchus punctatus* | | | | | | | | | | + | | | | | + | + | | | | | + | + | + | + | | | | |
| | *Parablennius yatabei* | | | + | | + | | + | | | | | | | + | | | | | + | + | + | + | + | | + | + | + | + |
| | *Petroscirtes breviceps* | | | | | | | | | | | | | | | | | | | | | | | | | | | | + |
| Callionymidae | *Repomucenus valenciennei* | | | | | | | | | | + | | | | | | | | | | | | | | | | | | |
| | *Repomucenus* spp. | | | | + | | | | | | | | | | | | | | | | | | | | | | | | |

Table S2. Continued.

| Family | Scientific Name | Aka | | | | | | Tama | | | | Miya | | | | | | Takatsu | | | | | | Sendai | | | | | |
|---|---|---|---|---|---|---|---|---|---|---|---|---|---|---|---|---|---|---|---|---|---|---|---|---|---|---|---|---|---|
| | | HT | LT | L1 | L2 | R1 | R2 | HT | LT | L2 | R1 | HT | LT | L1 | L2 | R1 | R2 | HT | LT | L1 | L2 | R1 | R2 | HT | LT | L1 | L2 | R1 | R2 |
| Odontobutidae | *Odontobutis hikimius* ‡ | | | | | | | | | | | | | | | | | | + | | | | | | | | | | |
| | *Odontobutis obscura* | | | | | | | | | | | | | | | | | + | + | + | + | + | + | | | | | | |
| Eleotridae | *Eleotris oxycephala* | | | | | | | | | | | | | | | | | + | + | + | + | + | + | | | | | | |
| Gobiidae | *Acanthogobius flavimanus* | + | + | | | | | | + | + | + | + | + | + | + | + | + | | | + | | | + | | + | + | | | |
| | *Acanthogobius lactipes* | | | | | | | | | | | | + | | | | | | | | | + | + | | | | | | |
| | *Acentrogobius* spp. | | | | | | | | | | | | | | | | | | | | | | | + | | | | | |
| | *Bathygobius hongkongensis* | | | | | | | | | | | | | | | | | | | | | | | | | + | | | |
| | *Chaenogobius annularis* | | | + | | | + | | | | | | | | | | | | | | | | | | | | | | |
| | *Chaenogobius gulosus* | | | + | | | | | | | | | | | | | | | | | | | | | | | | | |
| | *Eutaeniichthys gilli* ‡ | | | | | | | | | | | | | | | | | | | | | | | | | + | + | | |
| | *Favonigobius gymnauchen* | | | | | | | | | | | | | | | + | | | | | | | | | | | | | |
| | *Glossogobius olivaceus* | | | | | | | | | | | | | | | | | | | | | | | + | | | | | |
| | *Gymnogobius breunigii* | | | | | | | | + | | | + | + | + | + | + | + | | | | | | | | | | | | |
| | *Gymnogobius castaneus* ‡ | + | + | | | + | | | | | | | | | | | | | | | | | | | | | | | |
| | *Gymnogobius heptacanthus* | | | | | | | | | | | | | + | | | | | | | | | | | | | | | |
| | *Gymnogobius opperiens* | | + | | | | | | | | | | | | | | | | | | | | | | | | | | |
| | *Gymnogobius petschiliensis* | + | | | | | | | | | | | | + | | | | + | + | + | + | + | + | | | | | | |
| | *Gymnogobius scrobiculatus* ‡ | | | | | | | | | | | + | + | + | | | | | | | | | + | | | | | | |
| | *Gymnogobius urotaenia* | + | + | + | | | | | | | | | | + | + | | + | | + | + | | + | | | | | | | |
| | *Istigobius campbelli* | | | | | | | | | | | | | | | | | | | | | | | | | | | | + |
| | *Luciogobius guttatus* | + | | + | | | | | | | | + | + | + | | + | + | + | + | + | | + | + | | | | | | |
| | *Luciogobius pallidus* | | | | | | | | | | | | | | | | | + | + | + | | + | + | | | | | | |
| | *Luciogobius platycephalus* | | | | | | | | | | | | | + | + | + | + | | | | | | | | | | | | |
| | *Redigobius bikolanus* | | | | | | | | | | | | | | | | | | | + | + | | | | | | | | |
| | *Rhinogobius similis* | | | | | | | | | | | + | + | + | + | | | + | + | + | + | + | + | | | + | + | | |
| | *Rhinogobius* spp. | + | + | | | | | | | | | + | | + | + | | | + | + | + | + | + | + | + | + | | + | | |
| | *Taenioides snyderi* | | | | | | | | | | | | | | | | | | | | | | | | + | + | + | | |
| | *Tridentiger trigonocephalus* | | | | | | | | + | | | | | | | | | | | | | | | + | | + | + | | |
| | *Tridentiger* spp. | + | + | | | | | | | | | + | + | + | | + | + | + | + | + | + | + | + | + | + | + | + | | |
| Ptereleotridae | *Parioglossus dotui* | | | | | | | | | | | | | | | | | | | | | | | | + | | | | |
| Scatophagidae | *Scatophagus argus* | | | | | | | | | | | | | | | | | | | | | | | + | + | + | + | + | |
| Siganidae | *Siganus fuscescens* | | | | | | | | | | | | | + | + | | | | | + | + | + | + | | | + | | + | |
| Acanthuridae | *Prionurus scalprum* | | | | | | | | | | | | | | | | | | | | | | | + | | | | | |
| Sphyraenidae | *Sphyraena japonica* | | | | | | | | | | | | | | | | | | | | | | | + | | + | | | |
| | *Sphyraena pinguis* | | | | + | | | + | | | | | | | | | | | | + | + | + | + | + | | + | + | + | + |
| | *Sphyraena obtusata* | | | | | | | | | | | | | | | | | | | | | | | | | | | + | |
| Scombridae | *Auxis* spp. | | | | | | | | | | | | | | | | | | | | | | | + | | | | | |
| | *Scomber* spp. | | | + | + | + | | + | | + | | | | | | | | | | + | | | + | | | | | | |
| | *Scomberomorus niphonius* | | | | | | | | | | | | | | | | | | | | | | + | | | | | | |
| Paralichthyidae | *Paralichthys olivaceus* | + | | + | + | + | + | | | | | | | + | + | | + | | | | | | | | | | | | |
| Pleuronectidae | *Kareius bicoloratus* | | | | | | | | + | | | + | + | + | + | + | + | | | + | | | | | | | | | |
| | *Platichthys stellatus* | | | | | + | | | | | | | | | | | | | | | | | | | | | | | |
| | *Pseudopleuronectes yokohamae* | | | | | | | | | | | | | + | + | | | | | | | | | | | | | | |
| Soleidae | *Heteromycteris japonicus* | | | | + | | | | | | | | | | | | | | | | | | | | | | | | |
| Cynoglossidae | *Paraplagusia japonica* | | | | | | | | | | | | | | | | + | | | | | | | | | | | | |
| Monacanthidae | *Rudarius ercodes* | | | | | | + | | | | | | | | | | | | | | | | | | | | | | |
| | *Stephanolepis cirrhifer* | | | | | | | | | | | | | | | | | | | + | + | | | | | | | + | + |
| Ostraciidae | *Ostracion* spp. | | | | | | | | | | | | | | | | | | | | | | | | | | | | + |
| Tetraodontidae | *Canthigaster rivulata* | | | | | | | | | | | | | | | | | | | | | | | | | | | + | + |
| | *Takifugu* spp. | + | + | + | + | + | + | | | + | | + | + | + | + | + | + | + | | + | + | + | + | + | + | + | + | + | + |
| Number of Species | | 37 | 34 | 25 | 20 | 25 | 19 | 12 | 12 | 10 | 13 | 34 | 32 | 51 | 39 | 17 | 29 | 32 | 33 | 59 | 50 | 49 | 51 | 41 | 27 | 43 | 33 | 39 | 42 |
| Station | | HT | LT | L1 | L2 | R1 | R2 | HT | LT | L2 | R1 | HT | LT | L1 | L2 | R1 | R2 | HT | LT | L1 | L2 | R1 | R2 | HT | LT | L1 | L2 | R1 | R2 |
| River | | Aka | | | | | | Tama | | | | Miya | | | | | | Takatsu | | | | | | Sendai | | | | | |
| Number of Species (Total) | | 64 | | | | | | 25 | | | | 72 | | | | | | 81 | | | | | | 94 | | | | | |
| Number of Endangered Species (IUCN)† | | 2 | | | | | | 1 | | | | 4 | | | | | | 5 | | | | | | 3 | | | | | |
| Number of Endangered Species (Japan)‡ | | 6 | | | | | | 0 | | | | 7 | | | | | | 8 | | | | | | 2 | | | | | |
